# Genetic manipulation of insulin/insulin-like growth factor signaling pathway activity has sex-biased effects on *Drosophila* body size

**DOI:** 10.1101/2020.09.04.283382

**Authors:** Jason W. Millington, George P. Brownrigg, Paige J. Basner-Collins, Ziwei Sun, Elizabeth J. Rideout

**Affiliations:** Department of Cellular and Physiological Sciences, Life Sciences Institute, The University of British Columbia, Vancouver, BC, Canada, V6T 1Z3

**Keywords:** *Drosophila*, sex, insulin pathway, body size, genetics

## Abstract

In *Drosophila*, female body size is approximately 30% larger than male body size due to an increased rate of larval growth. While the mechanisms that control this sex difference in body size remain incompletely understood, recent studies suggest that the insulin/insulin-like growth factor signaling pathway (IIS) plays a role in the sex-specific regulation of growth during development. In larvae, IIS activity differs between the sexes, and there is evidence of sex-specific regulation of IIS ligands. Yet, we lack knowledge of how changes to IIS activity impact growth in each sex, as the majority of studies on IIS and body size use single- or mixed-sex groups of larvae and/or adult flies. The goal of our current study was to clarify the requirement for IIS activity in each sex during the larval growth period. To achieve this goal we used established genetic approaches to enhance, or inhibit, IIS activity, and quantified body size in male and female larvae. Overall, genotypes that inhibited IIS activity caused a female-biased decrease in body size, whereas genotypes that augmented IIS activity caused a male-specific increase in body size. This data extends our current understanding of larval growth by showing that most changes to IIS pathway activity have sex-biased effects on body size, and highlights the importance of analyzing data by sex in larval growth studies.

## INTRODUCTION

Over the past two decades, the *Drosophila* larva has emerged as an important model to study the regulation of growth during development. One important factor that affects body size in most *Drosophila* species is whether the animal is male or female: female flies are typically larger than male flies (Alpatov et al., 1930; Pitnick et al., 1995; French et al., 1998; Huey et al., 2006; Testa et al., 2013; Okamoto et al., 2013; Rideout et al., 2015; Sawala and Gould, 2017; Millington et al., 2020; reviewed in Millington and Rideout, 2018). This increased body size is due to an increased rate of larval growth, as the duration of the larval growth period does not differ between the sexes (Testa et al., 2013; Okamoto et al., 2013; Sawala and Gould, 2017). While the precise molecular mechanisms underlying the male-female difference in body size remain incompletely understood, recent studies have revealed a key role for the insulin/insulin-like growth factor signaling pathway (IIS) in the sex-specific regulation of growth during development (Shingleton et al., 2005; Gronke et al., 2010; Testa et al., 2013; Rideout et al., 2015; Liao et al., 2020; Millington et al., 2020).

Normally, IIS activity is higher in female larvae than in age-matched males (Rideout et al 2015; Millington et al., 2020). Given that increased IIS activity is known to promote cell, tissue, and organismal growth (Grewal, 2009; Teleman, 2009), this suggests that elevated IIS activity is one reason that females have an increased rate of growth and a larger body size. Indeed, the sex difference in growth was abolished between male and female flies carrying a mutation that strongly reduced IIS activity (Testa et al., 2013), and between male and female larvae reared on diets that markedly decrease IIS activity (Rideout et al., 2015). In both cases, the sex difference in growth was eliminated by a female-biased decrease in body size (Testa et al., 2013; Rideout et al., 2015). While these findings suggest that IIS plays a role in the sex-specific regulation of growth during development, only one genetic combination was used to reduce IIS activity (Testa et al., 2013). Therefore, it remains unclear whether the sex-biased effect of reduced IIS activity on body size is a common feature of genotypes that alter IIS activity.

In the present study, we used multiple genetic approaches to either enhance or inhibit IIS activity, and monitored larval growth in males and females. While previous studies show that these genetic approaches effectively alter IIS activity, the body size effects in each sex remain unclear due to frequent use of mixed-sex or single-sex experimental groups, and a lack of statistical tests to detect sex-by-genotype interactions (Fernandez et al., 1995; Chen et al., 1996; Leevers et al., 1996; Böhni et al., 1999; Brogiolo et al., 2001; Cho et al., 2001; Rintelen et al., 2001; Ikeya et al., 2002; Britton et al., 2002; Rulifson et al., 2002; Zhang et al., 2009; Geminard et al., 2009; Gronke et al., 2010). Our systematic examination of IIS revealed most genetic manipulations that reduced IIS activity caused a female-biased reduction in body size. In contrast, most genetic manipulations that enhanced IIS activity increased male body size with no effect in females. Together, these findings provide additional genetic support for IIS as an important regulator of sex-specific growth in *Drosophila*.

## MATERIALS AND METHODS

### Data Availability

Original images of pupae are available upon request. Raw values for all data collected and displayed in this manuscript are available in Supplementary file 1. The authors affirm that all data necessary for confirming the conclusions of the article are present within the article, figures, tables, and Supplementary files.

### Fly husbandry

*Drosophila* growth medium consisted of: 20.5 g/L sucrose, 70.9 g/L D-glucose, 48.5 g/L cornmeal, 45.3 g/L yeast, 4.55 g/L agar, 0.5g CaCl_2_•2H_2_O, 0.5 g MgSO_4_•7H_2_O, 11.77 mL acid mix (propionic acid/phosphoric acid). Diet data was deposited under “Rideout Lab 2Y diet” in the *Drosophila* Dietary Composition Calculator (Lesperance and Broderick, 2020). Larvae were raised at a density of 50 animals per 10 mL food at 25°C, and sexed by gonad size. Adult flies were maintained at a density of twenty flies per vial in single-sex groups.

### Fly strains

The following fly strains from the Bloomington *Drosophila* Stock Center were used: *w^1118^* (#3605), *UAS-rpr* (#5823), *UAS-Imp-L2-RNAi* (#55855), *InR^E19^* (#9646), *InR^PZ^* (#11661), *Df(3R)Pi3K92EA* (#25900), *chico^1^* (#10738), *foxo^21^* (#80943), *foxo^25^* (#80944), *r4-GAL4* (fat body), *dilp2-GAL4* (IPCs). Additional fly strains include: *UAS-Kir2.1* (Baines et al., 2001), *dilp1*, *dilp3, dilp4, dilp5, dilp6^41^, dilp7, Df(3L)ilp2-3,5, Df(3L)ilp1-4,5* (Grönke et al., 2010)*, Sdr^1^* (Okamoto et al., 2013), *Pi3K92E^2H1^* (Halfar et al., 2001), *Pdk1^4^* (Rintelen et al., 2001), *Akt1^3^* (Stocker et al., 2002). All fly strains except *dilp6^41^* were backcrossed into a *w^1118^* background.

### Body size

Pupal length and width were determined using an automated detection and measurement system. Segmentation of the pupae for automated analysis was carried out using the “Marker-controlled Watershed” function included in the MorphoJ plugin (Klingenberg, 2011) in ImageJ (Schindelin et al., 2012; Rueden et al., 2017). Briefly, the original image containing the pupae was blurred using the “Gaussian blur” function. A selection of points marking the pupae was then created using the “Find Maxima” function. Next, a new image with the same dimension as the pupae was created, where the individual points were projected onto this original image using the “Draw” function. Then, we labelled each point using the “Connected Components Labeling” function in the MorphoJ plugin (Klingenberg, 2011). This image is now the marker image. Upon completion of the marker image, we used the “Morphological Filters” function in the MorphoJ package with the options “operation=Gradient element=Octagon radius =2” to generate a gradient image of the pupae. Using the “Marker-controlled Watershed” function with the gradient image as the input, and the marker image to identify regions of interest outlining the pupae, the width and length of the pupae were obtained by selecting “Fit ellipse” option under the “Set Measurements” menu in ImageJ. Once length and width were determined using this automated measurement system, pupal volume was calculated as previously described (Delanoue et al., 2010; Rideout et al., 2012, 2015; Marshall et al., 2012; Ghosh et al., 2014). To measure adult weight, 5-day-old virgin male and female flies were collected and weighed in groups of ten on an analytical balance.

### Statistical analysis and data presentation

GraphPad Prism (GraphPad Prism version 8.4.2 for Mac OS X) was used to perform all statistical tests and to prepare all graphs in this manuscript. Statistical tests are indicated in figure legends and all *p*-values are listed in Supplementary file 2.

## RESULTS

### Reduced IPC function causes a female-biased decrease in body size

In *Drosophila*, the insulin-producing cells (IPCs) located in the brain are an important source of IIS ligands called *Drosophila* insulin-like peptides (Dilps). In larvae, the IPCs synthesize and release Dilp1 (<u>FBgn0044051</u>), Dilp2 (<u>FBgn0036046</u>), Dilp3 (<u>FBgn0044050</u>), and Dilp5 (<u>FBgn0044048</u>) into the hemolymph (Brogiolo et al., 2001; Ikeya et al., 2002; Rulifson et al., 2002; Lee et al., 2008; Geminard et al., 2009). When circulating Dilps bind to the Insulin-like Receptor (InR; <u>FBgn0283499</u>) on the surface of target tissues, an intracellular signaling cascade is initiated which ultimately promotes cell, tissue, and organismal growth (Chen et al., 1996; Böhni et al., 1999; Poltilove et al., 2000; Britton et al., 2002; Werz et al., 2009; Almudi et al., 2013). The importance of the IPCs in regulating IIS activity and growth is illustrated by the fact that IPC ablation and silencing both reduce IIS activity and decrease overall body size (Rulifson et al., 2002; Geminard et al., 2009). Yet, the precise requirement for IPCs in regulating growth in each sex remains unclear, as past studies presented data from a mixed-sex population of larvae or reported effects in only a single sex (Rulifson et al., 2002; Geminard et al., 2009). Given that recent studies show that the sex of the IPCs contributes to the sex-specific regulation of larval growth (Sawala and Gould, 2017), we asked how the presence and function of the IPCs affected body size in each sex.

First, we ablated the IPCs by overexpressing proapoptotic gene *reaper* (*rpr*; <u>FBgn0011706</u>) with the IPC-specific GAL4 driver *dilp2-GAL4* (Brogiolo et al., 2001; Rulifson et al., 2002). This method eliminates the IPCs during development (Rulifson et al., 2002). To quantify body size, we measured pupal volume, as it is a sensitive readout for larval growth (Delanoue et al., 2010). In females, pupal volume was significantly lower in *dilp2>UAS-rpr* larvae compared with *dilp2>+* and *+>UAS-rpr* control larvae (Fig. 1A). In males, pupal volume was also significantly lower in *dilp2>UAS-rpr* larvae compared with control *dilp2>+* and *+>UAS-rpr* larvae (Fig. 1A); however, the magnitude of the decrease in body size was greater in females than in males (sex:genotype interaction *p*<0.0001; two-way ANOVA). Next, to determine how reduced IPC function affected body size in each sex, we overexpressed the inwardly-rectifying potassium channel *Kir2.1* (Baines et al., 2001) using *dilp2-GAL4*. This approach reduces Dilp secretion and lowers IIS activity in a mixed-sex group of larvae (Geminard et al., 2009). We found that pupal volume was significantly reduced in *dilp2>UAS-Kir2.1* females compared with *dilp2>+* and *+>UAS-Kir2.1* control females (Fig. 1B). In males, pupal volume was reduced in *dilp2>UAS-Kir2.1* larvae compared with *dilp2>+* and *+>UAS-Kir2.1* control larvae (Fig. 1B). Because the magnitude of the decrease in female body size was larger than the reduction in male body size (sex:genotype interaction *p*<0.0001; two-way ANOVA), this result indicates that inhibiting IPC function caused a female-biased reduction in growth. Together, these results identify a previously unrecognized sex-biased body size effect caused by manipulating IPC survival and function.

**Figure 1.**
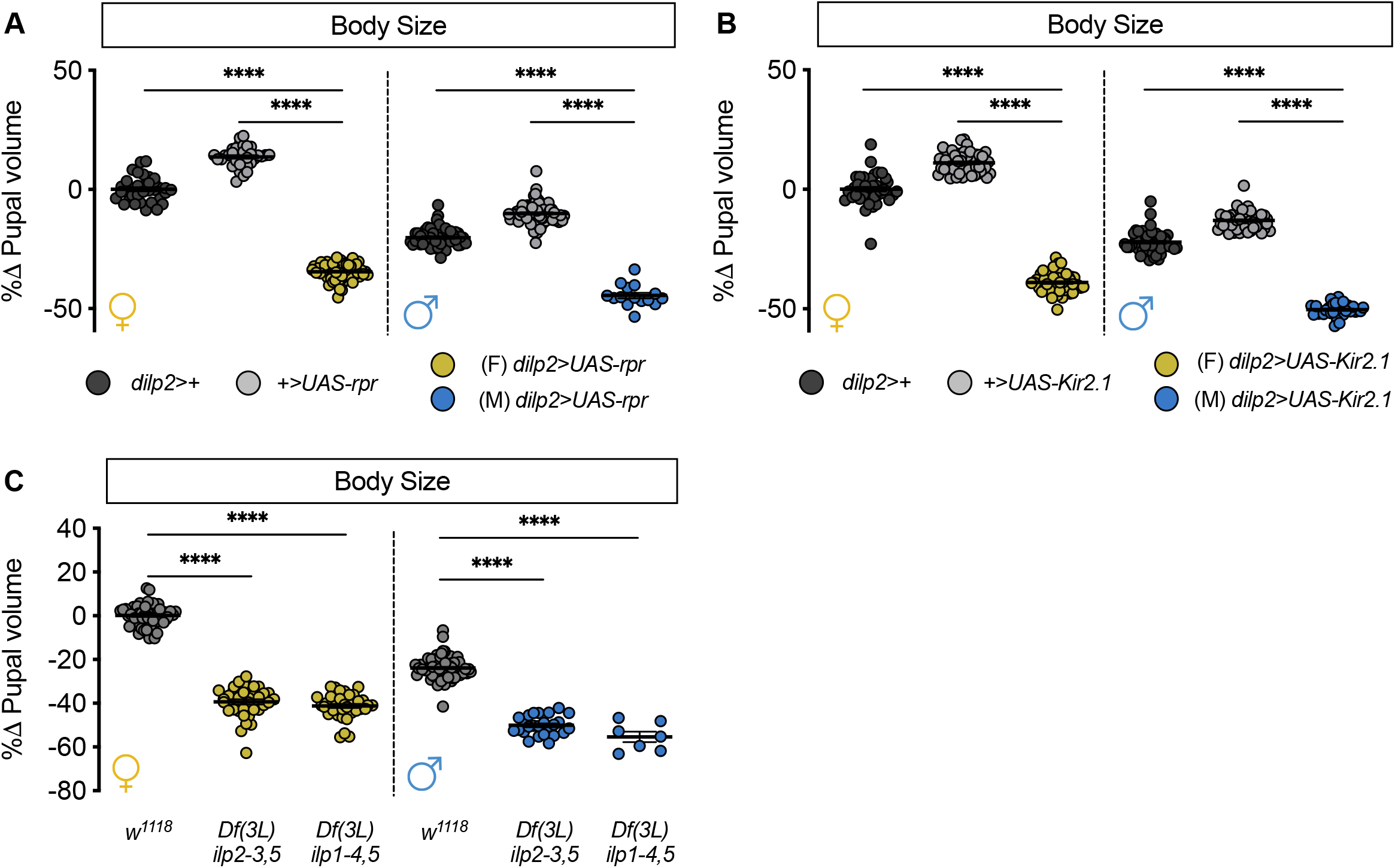
IPC ablation, loss of IPC function, and loss of IPC-derived Dilp ligands all cause a female-biased decrease in growth. (A) Pupal volume was significantly reduced in *dilp2>UAS-rpr* females and males compared to both *dilp2>+* and *+>UAS-rpr* controls (*p*<0.0001 for all comparisons; two-way ANOVA followed by Tukey HSD test). The magnitude of the reduction in pupal volume was greater in females (sex:genotype interaction *p*<0.0001; two-way ANOVA). *n* = 15-71 pupae. (B) Pupal volume was significantly reduced in *dilp2>UAS-Kir2.1* females and males compared to both *dilp2>+* and *+>UAS-Kir2.1* controls (*p*<0.0001 for all comparisons; two-way ANOVA followed by Tukey HSD test). The magnitude of the reduction in pupal volume was greater in females (sex:genotype interaction *p*<0.0001; two-way ANOVA followed by Tukey HSD test). *n* = 31-53 pupae. (C) Pupal volume was significantly reduced in *Df(3L)ilp2-3,5* homozygous females and males compared with sex-matched *w^1118^* controls (*p*<0.0001 for all comparisons; two-way ANOVA followed by Tukey HSD test). Similarly, *Df(3L)ilp1-4,5* homozygous females and males were significantly smaller than *w^1118^* control females and males (*p*<0.0001 for all comparisons; two-way ANOVA followed by Tukey HSD test). The magnitude of the reduction in body size for both *Df(3L)ilp2-3,5* and *Df(3L)ilp1-4,5* was significantly larger in females than in males (sex:genotype interaction *p*<0.0001 for both genotypes; two-way ANOVA followed by Tukey HSD test). *n* = 7-74 pupae. **** indicates *p*<0.0001; error bars indicate SEM. For all panels, females are shown on the left-hand side of the graph and males are shown on the right-hand side.

### Loss of IPC-derived Dilps causes a female-biased reduction in body size

Given that the larval IPCs produce Dilp1, Dilp2, Dilp3, and Dilp5 (Brogiolo et al., 2001; Ikeya et al., 2002; Rulifson et al., 2002; Lee et al., 2008; Geminard et al., 2009), we tested whether the loss of some (*Df(3L)ilp2-3,5*), or all (*Df(3L)ilp1-4,5*), of the IPC-derived Dilps affected larval growth in males and females. While a previous study reported how loss of all IPC-derived *dilp* genes affected adult weight, data from both sexes was not available for all genotypes (Gronke et al., 2010). In females, pupal volume was significantly smaller in *Df(3L)ilp2-3,5* larvae compared with *w^1118^* control larvae (Fig. 1C). In males, body size was also significantly reduced in *Df(3L)ilp2-3,5* homozygous larvae compared with *w^1118^* controls (Fig. 1C); however, the decrease in body size was significantly greater in females than in males (sex:genotype interaction *p*<0.0001; two-way ANOVA). When we measured body size in males and females lacking all IPC-derived Dilps (*Df(3L)ilp1-4,5), we* reproduced the female-biased reduction in body size (Fig. 1C; sex:genotype interaction *p*<0.0001; two-way ANOVA). This reveals a previously unrecognized sex-biased body size effect arising from loss of some, or all, IPC-derived Dilps.

### Loss of individual *dilp* genes causes a female-specific decrease in body size

While Dilp1, Dilp2, Dilp3 and Dilp5 are all produced by the IPCs, previous studies have uncovered significant differences in regulation, secretion, and phenotypic effects of these IPC-derived Dilps (Brogiolo et al., 2001; Zhang et al., 2009; Okamoto et al., 2009; Grönke et al., 2010; Cognigni et al., 2011; Stafford et al., 2012; Bai et al., 2012; Linneweber et al., 2014; Cong et al., 2015; Liu et al., 2016; Nässel & Vanden Broeck, 2016; Post et al., 2018, 2019; Semaniuk et al., 2018; Ugrankar et al., 2018; Brown et al., 2020). We therefore wanted to determine the individual contributions of IPC-derived Dilps to body size in each sex. Further, given that there are non-IPC-derived Dilps that regulate diverse aspects of physiology and behaviour (*dilp4*, <u>FBgn0044049</u>; *dilp6*, <u>FBgn0044047</u>; and *dilp7*, <u>FBgn0044046</u>) (Gronke et al., 2010; Castellanos et al., 2013; Garner et al., 2018), we wanted to determine the requirement for these additional Dilps in regulating larval growth in each sex. While a previous study measured adult weight as a read-out for body size in *dilp* mutants (Gronke et al 2010), we measured pupal volume to ensure changes to adult weight were not due to altered gonad size (Green and Extavour, 2014). We found that pupal volume was significantly smaller in female larvae carrying null mutations in *dilp1, dilp3, dilp4, dilp5*, and *dilp7* compared with *w^1118^* control females (Fig. 2A). This data aligns well with findings from two recent studies showing a female-specific decrease in larval growth caused by loss of *dilp2* (Liao et al., 2020; Millington et al., 2020). In contrast to most *dilp* mutants; however, there was no significant difference in pupal volume between homozygous *y,w,dilp6*^41^ female larvae and control *y,w* females (Fig. 2B). In males, pupal volume was not significantly different between *dilp1, dilp3, dilp4, dilp5*, and *dilp7* mutant larvae and *w^1118^* controls (Fig. 2C); however, pupal volume was significantly reduced in *y,w,dilp6^41^* larvae compared with *y,w* controls (Fig. 2D). Together, these results extend our current understanding of larval growth by revealing sex-specific requirements for all individual *dilp* genes in regulating body size.

**Figure 2.**
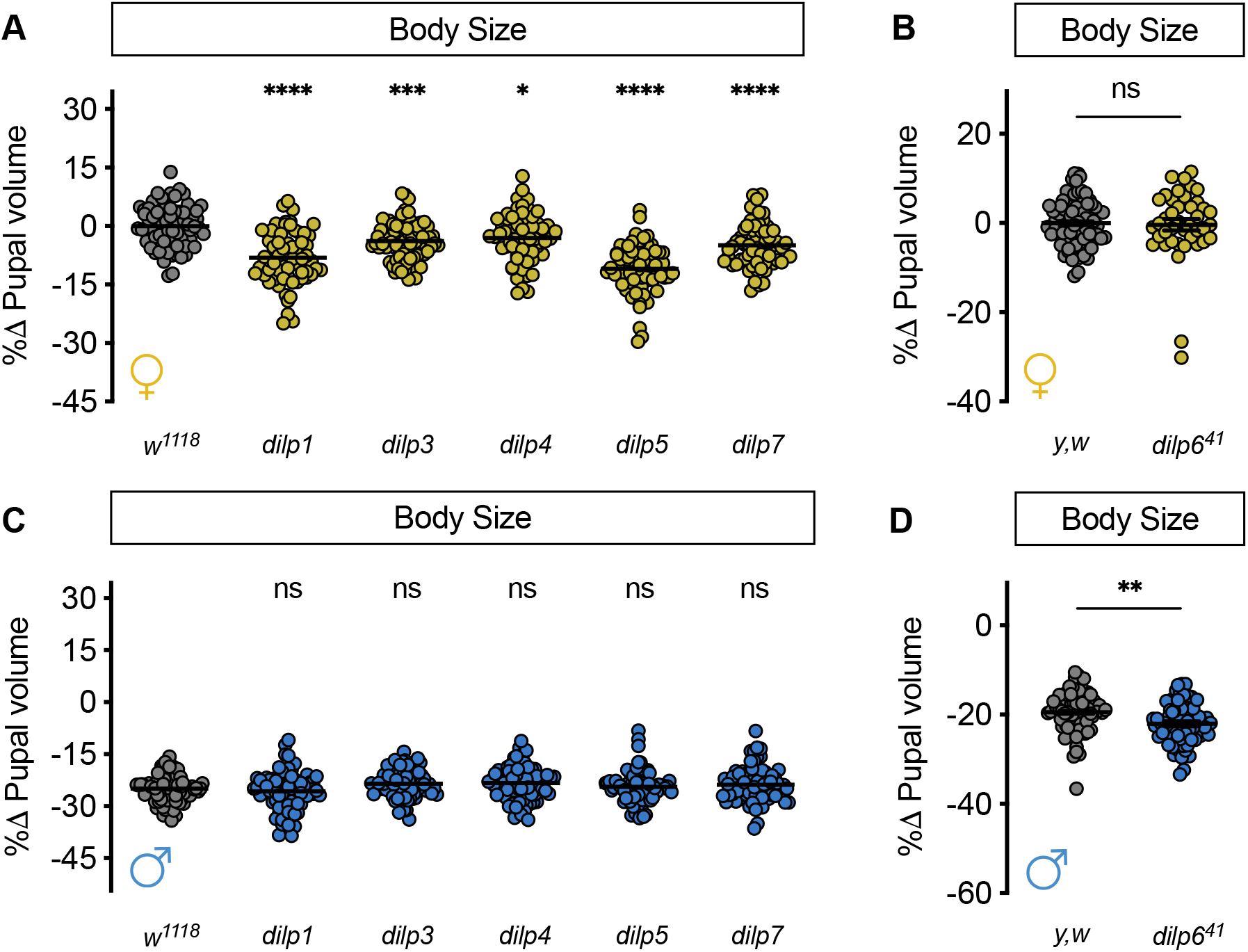
Loss of individual *dilp* genes causes sex-biased effects on growth. (A) In females, pupal volume was significantly reduced compared with *w^1118^* controls in larvae carrying individual mutations in each of the following genes: *dilp1, dilp3, dilp4, dilp5*, and *dilp7* (*p*<0.0001, *p* = 0.0003, *p* = 0.0136, *p*<0.0001, and *p*<0.0001, respectively; one-way ANOVA followed by Dunnett’s multiple comparison test). *n* = 59-74 pupae. (B) Pupal volume was not significantly different between *y,w* control female larvae and *dilp6^41^* mutant females (*p* = 0.7634, Student’s *t* test). *n* = 41-74 pupae. (C) In males, pupal volume was not significantly reduced compared with *w^1118^* controls in larvae carrying individual mutations in each of the following genes: *dilp1, dilp3, dilp4, dilp5*, and *dilp7 (p* = 0.7388, *p* = 0.2779, *p* = 0.1977, *p* = 0.9535, and *p* = 0.4526, respectively; one-way ANOVA followed by Dunnett’s multiple comparison test). *n* = 66-79 pupae. (D) Pupal volume was significantly reduced in male *dilp6^41^* larvae compared with *y,w* control males (*p* = 0.0017, Student’s *t* test). *n* = 64-70 pupae. * indicates *p*<0.05; ** indicates *p*<0.01; *** indicates *p*<0.001; **** indicates *p*<0.0001; ns indicates not significant; error bars indicate SEM. Panels A and B display female data; panels C and D show male data.

### Loss of Dilp-binding factor Imp-L2 causes a male-specific increase in body size

Once released into the circulation, the Dilps associate with proteins that modulate their growth-promoting effects. For example, Dilp1, Dilp2, Dilp5 and Dilp6 form a high-affinity complex with fat body-derived *ecdysone-inducible gene 2 (Imp-L2*, <u>FBgn0001257</u>) and Convoluted/*Drosophila* Acid Labile Subunit (Conv/dALS; <u>FBgn0261269</u>) (Okamoto et al., 2013), whereas Dilp3 interacts with Secreted decoy receptor of Insulin-like Receptor (Sdr; <u>FBgn0038279</u>) (Okamoto et al., 2013). Binding of the Imp-L2/dALS complex to individual Dilps likely reduces Dilp binding to InR, as reduced fat body levels of either Imp-L2 or dALS augment IIS activity and increase body size (Arquier et al., 2008; Honegger et al., 2008; Alic et al., 2011; Okamoto et al., 2013). Similarly, loss of Sdr increases IIS activity and increases body size (Okamoto et al., 2013). While the Sdr study reported that the magnitude of the increase in adult weight was equivalent in both sexes (Okamoto et al., 2013), which we confirm using pupal volume (Fig. 3A; sex:genotype interaction *p* = 0.5261; two-way ANOVA), it remains unclear how the Imp-L2/dALS complex affects growth in each sex. We found that in females pupal volume was not significantly different between larvae with fat body-specific overexpression of an *Imp-L2-RNAi* transgene (*r4>UAS-Imp-L2-RNAi*) and control *r4>+* and *+>UAS-Imp-L2-RNAi* larvae (Fig. 3B). In contrast, pupal volume was significantly larger in *r4>UAS-Imp-L2-RNAi* male larvae compared with *r4>+* and *+>UAS-Imp-L2-RNAi* control males (Fig. 3B). This result demonstrates a male-specific increase in larval growth caused by reduced *Imp-L2* (sex:genotype interaction *p*<0.0001; two-way ANOVA), revealing a previously unrecognized sex-specific effect of the Imp-L2/dALS complex on body size.

**Figure 3.**
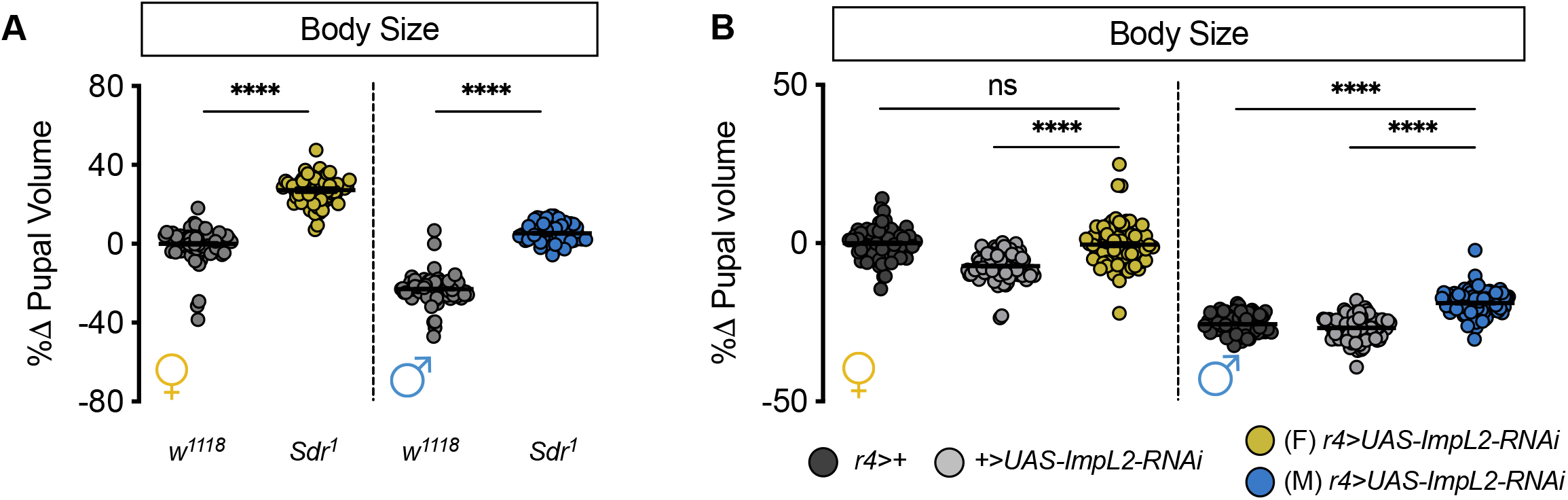
Fat body loss of Dilp-binding protein *Imp-L2* has sex-biased effects on growth. (A) Pupal volume was significantly increased in *Sdr^1^* mutant females and males compared with *w^1118^* control females and males (*p*<0.0001 for both sexes; two-way ANOVA followed by Tukey HSD test). There was no sex difference in the magnitude of the increase in body size (sex:genotype interaction *p* = 0.5261; two-way ANOVA followed by Tukey HSD test). *n* = 52-88 pupae. (B) In females, pupal volume was not significantly different between larvae with fat body-specific knockdown of *Imp-L2* (*r4>UAS-Imp-L2-RNAi*) compared with *r4>+* and *+>UAS-Imp-L2-RNAi* control larvae (*p* = 0.9948 and *p*<0.0001, respectively; two-way ANOVA followed by Tukey HSD test). In contrast, pupal volume was significantly larger in *r4>UAS-Imp-L2-RNAi* males compared with *r4>+* and *+>UAS-Imp-L2-RNAi* control males (*p*<0.0001 for both comparisons; two-way ANOVA followed by Tukey HSD test). The magnitude of the increase in pupal volume was higher in males than in females (sex:genotype interaction *p*<0.0001; two-way ANOVA). *n* = 70-92 pupae. **** indicates *p*<0.0001; ns indicates not significant; error bars indicate SEM. For all panels, females are shown on the left-hand side of the graph and males are shown on the right-hand side.

### Altered activity of the intracellular IIS pathway causes sex-biased and non-sex-specific effects on body size

In flies, IIS activity is stimulated by Dilp binding the InR on the surface of target cells (Fernandez et al., 1995; Chen et al., 1996). This Dilp-InR interaction induces receptor autophosphorylation and recruitment of adapter proteins such as Chico (<u>FBgn0024248</u>), the *Drosophila* homolog of mammalian insulin receptor substrate (Bohni et al., 1999; Poltilove et al., 2000; Werz et al., 2009). The recruitment and subsequent activation of the catalytic subunit of *Drosophila* phosphatidylinositol 3-kinase (*Pi3K92E*; <u>FBgn0015279</u>) increases the production of phosphatidylinositol (3,4,5)-trisphosphate (PIP3) at the plasma membrane (Leevers et al., 1996; Britton et al., 2002), which activates signaling proteins such as Phosphoinositide-dependent kinase 1 (Pdk1; <u>FBgn0020386</u>) and Akt1 (<u>FBgn0010379</u>) (Alessi et al., 1997). Both Pdk1 and Akt1 phosphorylate many downstream effectors to promote larval growth (Verdu et al., 1999; Cho et al., 2001; Rintelen et al., 2001). The importance of these intracellular IIS components in regulating growth during development is illustrated by studies showing that the loss, or reduced function, of most components decreases body size (Leevers et al., 1996; Chen et al., 1996; Böhni et al., 1999; Weinkove et al., 1999; Brogiolo et al., 2001; Rulifson et al., 2002; Zhang et al., 2009; Geminard et al., 2009; Grönke et al., 2010; Murillo-Maldonado et al., 2011). Yet, the majority of studies on the regulation of growth by intracellular IIS components were performed in a single- or mixed-sex population of larvae and/or adult flies, and lack testing for sex-by-genotype interactions (Fernandez et al., 1995; Chen et al., 1996; Leevers et al., 1996; Böhni et al., 1999; Brogiolo et al., 2001; Cho et al., 2001; Rintelen et al., 2001; Ikeya et al., 2002; Rulifson et al., 2002; Britton et al., 2002; Geminard et al., 2009; Zhang et al., 2009; Gronke et al., 2010). Given that recent studies have demonstrated the sex-specific regulation of IIS components such as Akt1 (Rideout et al., 2015), we investigated the requirement for these components in regulating larval growth in males and females. In line with previous results showing a female-biased decrease in adult weight in flies heterozygous for two hypomorphic *InR* alleles (Testa et al., 2013), we observed a female-biased pupal volume reduction in larvae carrying an additional combination of *InR* alleles (Fig. 4A; sex:genotype interaction *p*<0.0001; two-way ANOVA).

**Figure 4.**
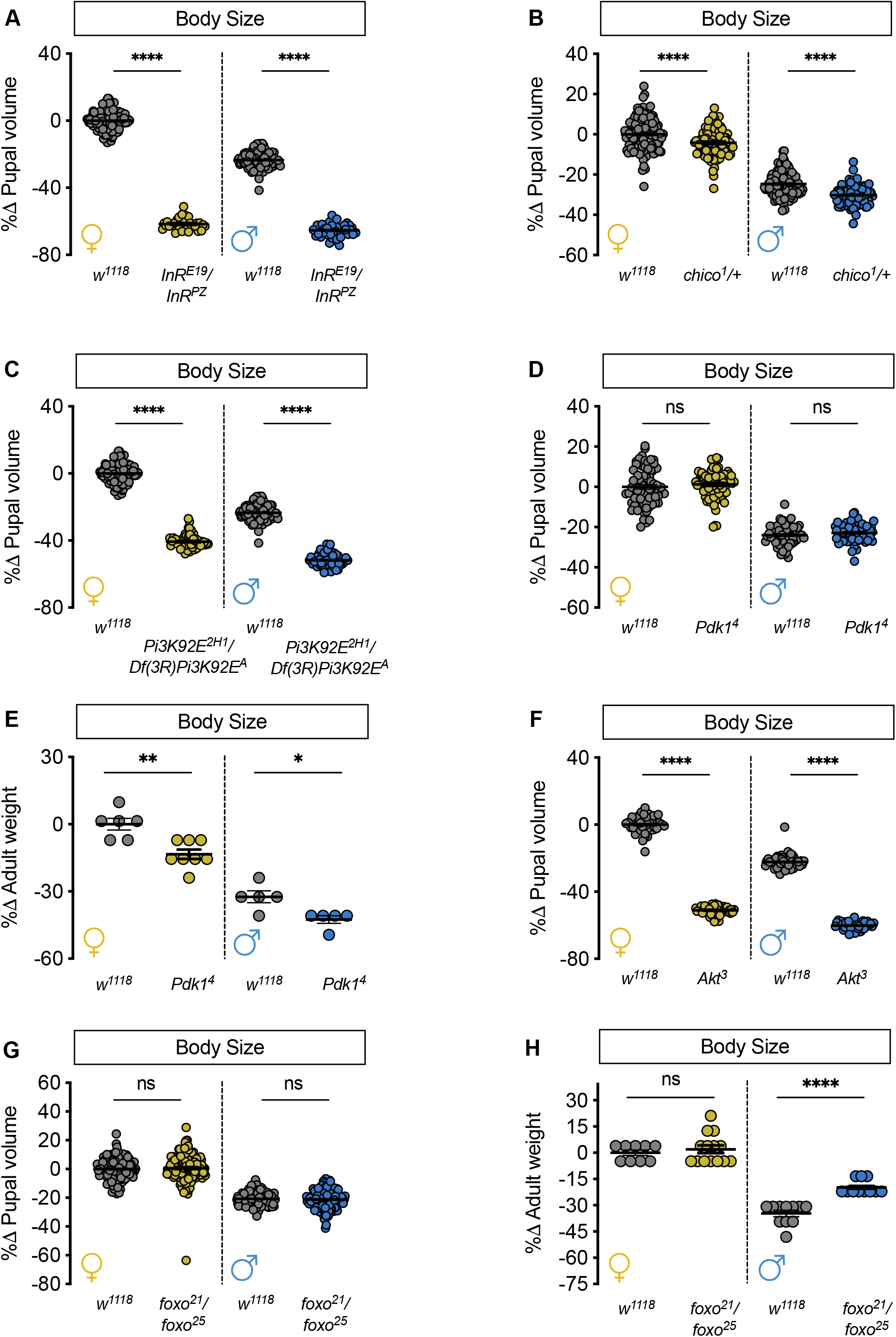
Both sex-biased and non-sex-biased effects on growth arise from loss of intracelllular IIS pathway components. (A) Pupal volume was significantly reduced in females and males heterozygous for two hypomorphic *InR* alleles (*InR^E19^/InR^PZ^*) compared with sex-matched *w^1118^* controls (*p*<0.0001 for both sexes; two-way ANOVA followed by Tukey HSD test). The magnitude of the decrease in larval body size was significantly higher in *InR^E19^/InR^PZ^* females than in *InR^E19^/InR^PZ^* males (sex:genotype interaction *p* = 0.0029; two-way ANOVA followed by Tukey HSD test). *n* = 32-133 pupae. (B) Pupal volume was significantly smaller in females and males heterozygous for a null *chico* allele (*chico^1^/+*) compared with sex-matched *w^1118^* controls (*p*<0.0001 for both females and males; two-way ANOVA followed by Tukey HSD test). The magnitude of the reduction in body size was not significantly different between females and males (sex:genotype interaction *p* = 0.1399; two-way ANOVA followed by Tukey HSD test). *n* = 93-133 pupae. (C) Pupal volume was significantly reduced in females and males heterozygous for a deficiency and hypomorphic allele of *Pi3K92E* (*Df(3R)Pi3K92E^A^/Pi3K92E^2H1^*) compared with sex-matched *w^1118^* controls (*p*<0.0001 for all comparisons in females and males; two-way ANOVA followed by Tukey HSD test). The magnitude of the reduction in body size was significantly larger in *Df(3R)Pi3K92E^A^/Pi3K92E^2H1^* females than in *Df(3R)Pi3K92E^A^/Pi3K92E^2H1^* males (sex:genotype interaction *p* = 0.0029; two-way ANOVA followed by Tukey HSD test). Note: the *Df(3R)Pi3K92E^A^/Pi3K92E^2H1^* pupae were collected and analyzed in parallel with the *InR^E19^/InR^PZ^* genotype, so the *w^1118^* control genotype data is shared between these experiments. *n* = 52-133 pupae. (D) Pupal volume was not significant different in either females or males homozygous for a hypomorphic *Pdk1* allele (*Pdk1^4^*) compared with *w^1118^* controls (*p* = 0.6739 and *p* = 0.7847, respectively; two-way ANOVA followed by Tukey HSD test). *n* = 61-84 pupae. (E) Adult weight was significantly reduced in *Pdk1^4^* females and males compared with *w^1118^* controls (*p* = 0.0017 and *p* = 0.0491 for females and males respectively; two-way ANOVA followed by Tukey HSD test). The magnitude of the reduction in body size was not significantly different between females and males (sex:genotype interaction *p* = 0.503; two-way ANOVA followed by Tukey HSD test). *n* = 5-8 biological replicates of ten adult flies. (F) Pupal volume was significantly reduced in females and males homozygous for a hypomorphic *Akt1* allele (*Akt1^3^*) compared with sex-matched *w^1118^* controls (*p*<0.0001 for both sexes; two-way ANOVA followed by Tukey HSD test). The magnitude of the decrease in body size in *Akt1^3^* larvae was significantly higher in females than in males (sex:genotype interaction *p*<0.0001; two-way ANOVA followed by Tukey HSD test). *n* = 44-60 pupae. (G) In females and males heterozygous for two hypomorphic alleles of foxo (*foxo^21^/foxo^25^*), pupal volume was not significantly different compared with sex-matched *w^1118^* controls (*p* = 0.8841 and 0.9646, respectively; two-way ANOVA followed by Tukey HSD test). *n* = 110-153 pupae. (H) In *foxo^21^/foxo^25^* females, adult weight was not significantly different compared with *w^1118^* controls (*p* = 0.8786; two-way ANOVA followed by Tukey HSD test). In males, adult weight was significantly higher in *foxo^21^/foxo^25^* flies compared with *w^1118^* control flies (*p*<0.0001; two-way ANOVA followed by Tukey HSD test). The magnitude of the increase in body size was greater in males than in females (sex:genotype interaction *p* = 0.0014; two-way ANOVA followed by Tukey HSD test). *n* = 5-8 biological replicates of 10 adult flies. * indicates *p*<0.05; ** indicates *p*<0.01; **** indicates *p*<0.0001; ns indicates not significant; error bars indicate SEM. For all panels, females are shown on the left-hand side of the graph and males are shown on the right-hand side.

To expand these findings beyond *InR, we* measured pupal volume in males and females with whole-body loss of individual intracellular IIS components. Given that we did not obtain viable pupae homozygous for a null mutation in *chico (chico^1^), we* measured pupal volume in *chico^1^/+* males and females. In *chico^1^/+* females, pupal volume was significantly reduced compared with control *w^1118^* larvae (Fig. 4B). In *chico^1^/+* males, pupal volume was reduced compared with control *w^1118^* larvae (Fig. 4B). Given that the magnitude of the reduction in pupal volume was similar in males and females (sex:genotype interaction *p* = 0.1399; two-way ANOVA), reduced *chico* did not cause a sex-biased effect on larval growth. In females heterozygous for two mutant alleles of *Pi3K92E (Df(3R)Pi3K92E^A^/Pi3K92E^2H1^*), pupal volume was significantly reduced compared with control *w^1118^* larvae (Fig. 4C). In *Df(3R)Pi3K92E^A^/Pi3K92E^2H1^* males, we observed a significant reduction in pupal volume (Fig. 4C); however, the magnitude of the decrease in body size was larger in females compared with males (sex:genotype interaction *p*<0.0001; two-way ANOVA). This indicates that loss of Pi3K92E caused a female-biased decrease in larval growth. Next, we examined body size in males and females homozygous for a hypomorphic allele of *Pdk1 (Pdk1^4^)*. We observed no effect on pupal volume in either sex in *Pdk1^4^* homozygotes (Fig. 4D); however, when we measured adult weight we found an equivalent body size reduction in *Pdk1^4^* males and females compared with sex-matched control *w^1118^* flies (Fig. 4E; sex:genotype interaction *p* = 0.5030; two-way ANOVA). This suggests that reduced *Pdk1* did not cause a sex-biased reduction in larval growth. One important target of *Pdk1* is the serine/threonine kinase Akt1. In females homozygous for a hypomorphic allele of *Akt1* (*Akt1^3^*), pupal volume was significantly reduced compared with control *w^1118^* larvae (Fig. 4F). In *Akt1^3^* males, we observed a significant reduction in body size compared with control *w^1118^* larvae (Fig. 4F). Given that the magnitude of the decrease in body size was larger in females than in males (sex:genotype interaction *p*<0.0001; two-way ANOVA), this indicates that loss of Akt1 caused a female-biased effect on larval growth. Together, these findings identify previously unrecognized sex-biased body size effects of reduced *Pi3K92E* and *Akt1*.

One downstream target of IIS that contributes to the regulation of growth is transcription factor *forkhead box, sub-group O* (*foxo*; <u>FBgn0038197</u>). When IIS activity is high, Akt1 phosphorylates Foxo to prevent Foxo from translocating to the nucleus (Puig et al., 2003). Given that Foxo positively regulates mRNA levels of many genes that are involved in growth repression and catabolism (Zinke et al., 2002; Junger et al., 2003; Kramer et al., 2003; Slack et al., 2011; Alic et al., 2011), elevated IIS activity promotes growth in part by inhibiting Foxo (Junger et al., 2003; Kramer et al., 2003). Because previous studies show increased Foxo nuclear localization and elevated Foxo target gene expression in males (Rideout et al., 2015; Millington et al., 2020), we examined how Foxo contributes to larval growth in each sex by measuring body size in females and males heterozygous for two hypomorphic *foxo* alleles (*foxo^21^/foxo^25^*). In *foxo^21^/foxo^25^* females and males, pupal volume was not significantly different from sex-matched *w^1118^* control larvae (Fig. 4G). In adult females, body weight was not significantly different between *foxo^21^/foxo^25^* mutants and control *w^1118^* flies (Fig. 4H); however, *foxo^21^/foxo^25^* adult males were significantly heavier than control *w^1118^* males (Fig. 4H). Because we observed a male-specific increase in body size (sex:genotype interaction *p* = 0.0014; two-way ANOVA), our data suggests that Foxo function normally contributes to the smaller body size of males. This reveals a previously unrecognized sex-specific role for Foxo in regulating body size.

## DISCUSSION

Many studies have demonstrated an important role for IIS in promoting cell, tissue, and organismal growth in response to nutrient input (Fernandez et al., 1995; Chen et al., 1996; Böhni et al., 1999; Britton et al., 2002; Grewal, 2009; Teleman, 2009). More recently, studies suggest that IIS also plays a role in the sex-specific regulation of larval growth (Testa et al., 2013; Rideout et al., 2015; Millington et al., 2020). However, the links between IIS and the sex-specific regulation of growth were made based on a limited number of genotypes that affected IIS activity. The goal of our current study was to determine whether the sex-biased larval growth effects observed in previous studies represent a common feature of genotypes that affect IIS activity. Overall, we found that the loss of most positive regulators of IIS activity caused a female-biased reduction in body size. On the other hand, loss of genes that normally repress IIS activity caused a male-specific increase in body size. Thus, most changes to IIS activity cause sex-biased, or sex-specific, effects on larval growth (summarized in Table 1), highlighting the importance of collecting and analyzing data from both sexes separately in studies that manipulate IIS activity and/or examine IIS-responsive phenotypes (*e.g*., lifespan, immunity).

**Table 1.**
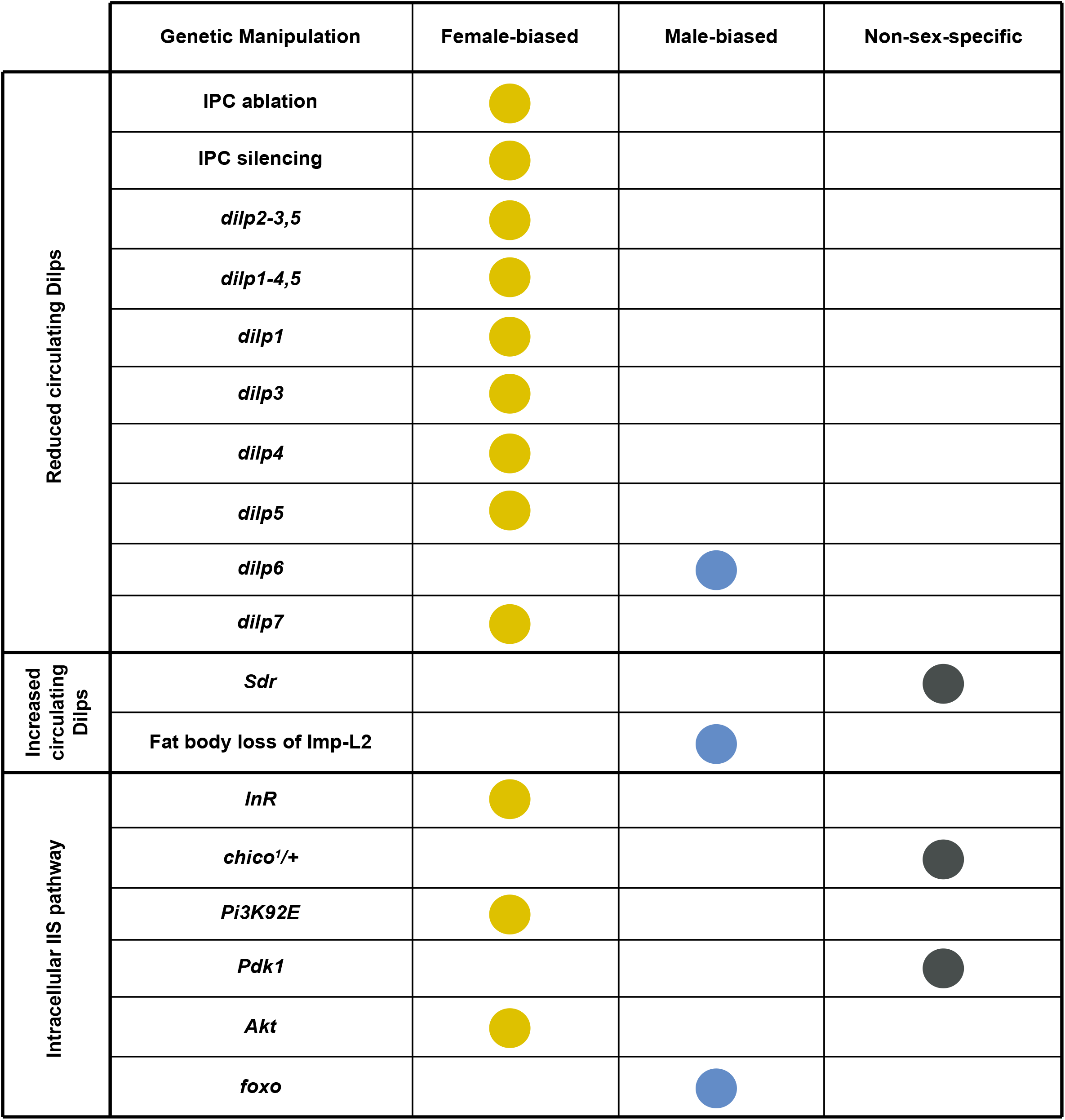
Summary of sex-biased effects of IIS pathway manipulations on body size.

One important outcome from our study was to provide additional genetic support for IIS as an important regulator of the sex difference in larval growth. Data implicating IIS in the sex-specific regulation of body size first emerged from a detailed examination of the rate and duration of larval growth in both sexes (Testa et al., 2013). In this study, the authors reported a female-biased growth reduction in larvae with decreased InR function (Testa et al 2013). A subsequent study extended this finding by uncovering a sex difference in IIS activity: late third-instar female larvae had higher IIS activity than age-matched males (Rideout et al., 2015). The reasons for this increased IIS activity remain incompletely understood; however, Dilp2 secretion from the IPCs was higher in female larvae than in males (Rideout et al., 2015). Given that Dilp2 overexpression is known to augment IIS activity and enhance body size (Ikeya et al., 2002; Geminard et al., 2009), these findings suggest a model in which high levels of circulating Dilp2 (and possibly other Dilps) are required in females to achieve and maintain increased IIS activity and a larger body size. In males, lower circulating levels of Dilp2 lead to reduced IIS activity and a smaller body size. If this model is accurate, we predict that female body size will be more sensitive to genetic manipulations that reduce Dilp ligands and/or IIS activity. Previous studies provided early support for this model by demonstrating a female-biased reduction in growth due to strong *InR* inhibition and *dilp2* loss (Testa et al., 2013; Liao et al., 2020; Millington et al., 2020). Now, we provide strong genetic support for this model using multiple genetic manipulations to reduce IIS activity, confirming that *Drosophila* females depend on high levels of IIS activity to promote increased body size. One potential reason for this high level of IIS activity in females is to ensure successful reproduction, as IIS activity in females regulates germline stem cell divisions, ovariole number, and egg production (LaFever and Drummond-Barbosa, 2005; Hsu et al., 2008; Hsu and Drummond-Barbosa, 2009; Gronke et al., 2010; Extavour and Green, 2014). Unfortunately, this elevated level of IIS activity shortens lifespan, revealing an important IIS-mediated tradeoff between fecundity and lifespan in females (Broughton et al., 2005).

A second prediction of this model is that augmenting either circulating Dilp levels or IIS activity will enhance male body size. Indeed, we demonstrate that loss of *Imp-L2*, which increases free circulating Dilp levels (Arquier et al., 2008; Honegger et al., 2008; Alic et al., 2011; Okamoto et al., 2013), and loss of *foxo*, which mediates growth repression associated with low IIS activity (Junger et al., 2003; Kramer et al., 2003), both cause a male-specific increase in body size. Together, these findings suggest that the smaller body size of male larvae is partly due to low IIS activity. While the reason for lower IIS activity in males remains unclear, studies show that altered IIS activity in either of the two main cell types within the testis compromises male fertility (Ueishi et al., 2009; McLeod et al., 2010; Amoyel et al., 2014; Amoyel et al., 2016). Future studies will therefore need to determine how males and females each maintain IIS activity within the range that maximizes fertility. In addition, it will be important to determine whether the female-biased phenotypic effects of lower IIS activity that we observe, and which are also widespread in aging and lifespan studies (Clancy et al., 2001; Holzenberger et al., 2003; Magwere et al., 2004; Van Heemst et al., 2005; Selman et al., 2008; Regan et al., 2016; Kane et al., 2018) extend to additional IIS-associated phenotypes (*e.g*., immunity and sleep) (DiAngelo et al., 2009; Cong et al., 2015; Roth et al., 2018; Suzawa et al., 2019; Brown et al., 2020).

Another important task for future studies will be to gain deeper insight into sex differences in the IPC function, as one study identified sex-specific Dilp2 secretion from the IPCs (Rideout et al., 2015). Indeed, recent studies have revealed the sex-specific regulation of one factor (*stunted*, <u>FBgn0014391</u>) that influences Dilp secretion from the IPCs (Millington et al., 2020), and female-specific phenotypic effects of another factor that influences IPC-derived Dilp expression (Woodling et al., 2020). Together, these studies suggest that sex differences in IPC function and circulating Dilp levels exist, and may arise from the combined effects of multiple regulatory mechanisms. Given that our knowledge of IPC function has recently expanded in a series of exciting studies (Meschi et al., 2019; Oh et al., 2019), more work will be needed to test whether these newly discovered modes of IPC regulation operate in both sexes. Further, it will be important to ascertain how sex differences in the IPCs are specified. One recent study showed that *Sex-lethal* (*Sxl*; <u>FBgn0264270</u>), a key regulator of female sexual development, acts in the IPCs to regulate the male-female difference in body size (Sawala and Gould, 2017). By studying how *Sxl* function alters IPC gene expression, activity, and connectivity, it will be possible to gain vital mechanistic insight into the sex-specific regulation of larval growth.

Beyond an improved understanding of sex differences in IPC function, it will be essential to study the sex-specific regulation of *dilp* genes and Dilp proteins, as we show female-specific effects on growth in larvae lacking individual *dilp* genes. While previous studies have reported female-biased effects of loss of *dilp2* (Liao et al 2020; Millington et al 2020), this is the first report of a female-specific role for *dilp1, dilp3, dilp4, dilp5*, and *dilp7* in promoting growth. Because loss of individual *dilp* genes reduced body size by ~10%, whereas loss of *InR* reduced body size by ~50%, we propose that increased levels of several Dilps contributes to the increased IIS activity and larger body size in females. While previous studies suggest that circulating Dilp2 levels are higher in female larvae (Rideout et al., 2015), it remains unclear whether other Dilps show similar sex-specific regulation. Interestingly, a recent study showed that in females there are more *dilp7*-positive cells than males due to programmed cell death in a subpopulation of male *dilp7*-positive cells (Castellanos et al., 2013; Garner et al., 2018). Given our finding that loss of *dilp7* causes a female-specific reduction in body size, it is possible that circulating Dilp7 levels also differ between the sexes. In the future, it will therefore be necessary to systematically analyze circulating levels of other Dilps in both sexes. Further, as our knowledge of how individual *dilp* genes affect larval development and physiology continues to grow, continued studies on the sex-specific regulation of *dilp* genes and Dilp proteins will be important to improve our understanding of male-female differences in larval growth, and to extend knowledge of sex differences in other IIS-associated traits.

In contrast to the female-biased effects of all genetic manipulations that reduced Dilp availability, we observed both sex-biased and non-sex-biased effects on body size in larvae with reduced function of key intracellular IIS components. For example, reduced InR, Pi3K92E, and Akt1 function caused a female-biased reduction in body size, whereas there was an equivalent reduction in male and female body size due to lower *chico* and *Pdk1* function. While the reasons for the lack of sex-biased effects of these two genes are unclear, one recent study showed that heterozygous loss of *chico* caused insulin hypersecretion (Sanaki et al., 2020). Given that hyperinsulinaemia contributes to insulin resistance, and that insulin resistance decreases *Drosophila* body size (Musselman et al., 2011, 2018; Pasco and Leopold, 2012), more studies will be needed to determine whether the smaller body size of *chico^1^/+* male and female larvae, and possibly *Pdk1* mutant larvae, can be attributed to insulin resistance. In fact, more knowledge of sex-specific tissue responses to insulin is urgently needed in flies, as studies in mice and humans have identified sex differences in insulin sensitivity (Macotela et al., 2009; Geer and Shen, 2009). Because *Drosophila* is an emerging model to understand the mechanisms underlying the development of insulin resistance (Musselman et al. 2011), this knowledge would help determine whether flies are a good model to investigate the sex-biased incidence of diseases associated with insulin resistance, such as the metabolic syndrome and type 2 diabetes (Mauvais-Jarvis, 2015).

## ACKNOWLEDGEMENTS

We would like to thank Chien Chao for developing the automated pupal volume analysis method. We would like to thank Dr. Linda Partridge for sharing the *dilp* mutant strains used in this study, Dr. Takashi Nishimura for sharing *Sdr^1^* flies, and Dr. Ernst Hafen for providing *Akt1^3^, Pdk1^4^*, and *Pi3K92E^2H1^* mutant strains. Stocks obtained from the Bloomington *Drosophila* Stock Center (NIH P40OD018537) were used in this study. We thank the TRiP at Harvard Medical School (NIH/NIGMS R01-GM084947) for providing transgenic RNAi fly stocks and/or plasmid vectors used in this study. We acknowledge critical resources and information provided by FlyBase (Thurmond, J. 2019, NAR); FlyBase is supported by a grant from the National Human Genome Research Institute at the U.S. National Institutes of Health (<u>U41 HG000739</u>) and by the British Medical Research Council (<u>MR/N030117/1</u>). Funding for this study was provided by grants to EJR from the Canadian Institutes for Health Research (PJT-153072), Natural Sciences and Engineering Research Council of Canada (NSERC, RGPIN-2016-04249), Michael Smith Foundation for Health Research (16876), and the Canadian Foundation for Innovation (JELF-34879). JWM was supported by a 4-year CELL Fellowship from UBC, and ZS was supported by an NSERC Undergraduate Student Research Award. We would like to acknowledge that our research takes place on the traditional, ancestral, and unceded territory of the Musqueam people; a privilege for which we are grateful.

